# Cross-omic Transcription Factors meta-analysis: an insight on TFs accessibility and expression correlation

**DOI:** 10.1101/2024.01.23.576789

**Authors:** Lorenzo Martini, Roberta Bardini, Alessandro Savino, Stefano Di Carlo

## Abstract

It is well-known how sequencing technologies propelled cellular biology research in recent years, giving an incredible insight into the basic mechanisms of cells. Single-cell RNA sequencing is at the front in this field, with Single-cell ATAC sequencing supporting it and becoming more popular. In this regard, multi-modal technologies play a crucial role, allowing the possibility to perform the mentioned sequencing modalities simultaneously on the same cells. Yet, there still needs to be a clear and dedicated way to analyze this multi-modal data. One of the current methods is to calculate the Gene Activity Matrix (GAM), which summarizes the accessibility of the genes at the genomic level, to have a more direct link with the transcriptomic data. However, this concept is not well-defined, and it is unclear how various accessible regions impact the expression of the genes. Moreover, the transcription process is highly regulated by the Transcription Factors that binds to the different DNA regions. Therefore, this work presents a continuation of the meta-analysis of Genomic-Annotated Gene Activity Matrix (GAGAM) contributions, aiming to investigate the correlation between the TFs expression and motif information in the different functional genomic regions to understand the different Transcription Factors (TFs) dynamics involved in different cell types.

## 1. Introduction

Next Generation Sequencing (NGS) technologies serve as the backbone for cutting-edge cellular biology research, offering a powerful tool to investigate fundamental cell mechanisms. These technologies, especially single-cell RNA sequencing (scRNA-seq) and single-cell assays for transposase-accessible chromatin sequencing (scATAC-seq), contribute significantly to studying cellular states with high resolution, a critical aspect for understanding cellular heterogeneity.

Widely utilized for profiling thousands of single-cell transcriptional profiles, scRNA-seq enables the investigation of cellular heterogeneity based on gene expression [1] [2] [3]. Simultaneously, the emerging popularity of scATAC-seq proves invaluable. This technology, by probing the entire genome and assessing accessible chromatin regions, offers complementary insights into gene regulation processes [4] and expression [5]. While the integration of scRNA-seq and scATAC-seq through multi-modal technologies is becoming crucial for understanding cell-related phenomena, the inherent differences in data types between the two technologies pose challenges to joint analysis [6] [7] [8]. Correlating the accessibility of a genomic region with gene expression is not straightforward due to the intricate machinery involved in transcriptional regulation. scRNA-seq datasets prioritize genes as prominent features, while scATAC-seq datasets consider genomic regions as features, making their integration challenging.

To bridge this gap, the GEne Activity (GA) concept is introduced, summarizing genomic accessibility information in a form where features are genes, allowing direct comparison with scRNA-seq matrices [9]. However, defining the relationship between accessible regions and genes remains unclear.

The Genomic-Annotated Gene Activity Matrix (GAGAM) approach [10] [11] proposes a promising solution, relying on a genomic model based on annotations to associate genomic regions with accessible genes. This approach constructs a Gene Activity Matrix (GAM) with contributions from different functional genomic regions (promoters, exons, and enhancers). Although GAGAM better models the gene regulatory landscape, it lacks representation of the complex gene regulation mechanisms [12], especially the involvement of Transcription Factors (TFs).

This work aims to address this gap by preliminary analyzing the correlation between TF expression and the accessibility of their motifs. Specifically, it explores differences in motif accessibility in promoter and enhancer regions, aiming to tailor TF information with GAGAM contributions nuancedly. This work is an extension of the previously published work documented in [13].

## 2. Background

To comprehend the proposed analysis, it is essential to introduce the fundamental technologies underpinning this work.

### 2.1 Single-cell sequencing technologies

A short overview of the scATAC-seq data organization aids in understanding the derived concept of Gene Activity (GA). scATAC-seq is a technology offering insights into the epigenomic state of cells by probing the entire genome. It utilizes the Tn5-transposase to identify regions where chromatin is open, and DNA sequences are accessible [14]. This technology enables the investigation not only of genes, as in scRNA-seq, but also of various functional elements such as enhancers and promoters scattered throughout the genome, crucial for gene regulation [15] [16].

While scRNA-seq data use genes as primary features, scATAC-seq data utilize peaks, i.e., short genomic regions described by their coordinates on chromosomes. This difference poses a significant challenge when correlating the two biological levels. One approach to overcome this hurdle is transforming peaks into gene-like data and comparing the two technologies. As the introduction mentions, GA serves as one such method [6].

However, current models for defining GA often oversimplify the relationship between a gene and the accessibility of its genomic region. Certain approaches, such as GeneScoring[17] and Signac [18], indiscriminately consider peak signals overlapping gene body regions without distinguishing between coding and non-coding regulatory elements. In contrast, Cicero [6] adopts a more structured approach, considering various regulatory regions but collapsing the gene region to a single base. These methods retain minimal biological information from raw scATAC-seq data, primarily related to gene coding regions, despite representing only a small percentage of the entire signal [19].

Beyond these simplistic models, other approaches aim to encompass more accessible genomic regions and their impact on overall GA. This work specifically employs GAGAM, utilizing curated genomic annotations to functionally label peaks and compute distinct contributions [10].

### 2.2 GAGAM

GAGAM uses information on various DNA regions, particularly exons and noncoding regions with regulatory roles, to improve the analysis of biological information from scATAC-seq data. This model-driven approach aims to support the study of cellular heterogeneity better. GAGAM lays the groundwork for a detailed investigation into the relationship between accessibility and expression in single-cell data. Its modular structure allows independent computation of contributions, facilitating specific and separate investigations, which is especially crucial when considering the role of regulatory regions with challenging relationships to gene expression. Algorithm 1 briefly overviews the workflow required to construct GAGAM contributions, starting from raw data, to provide the necessary background.

#### Algorithm 1

GAGAM construction

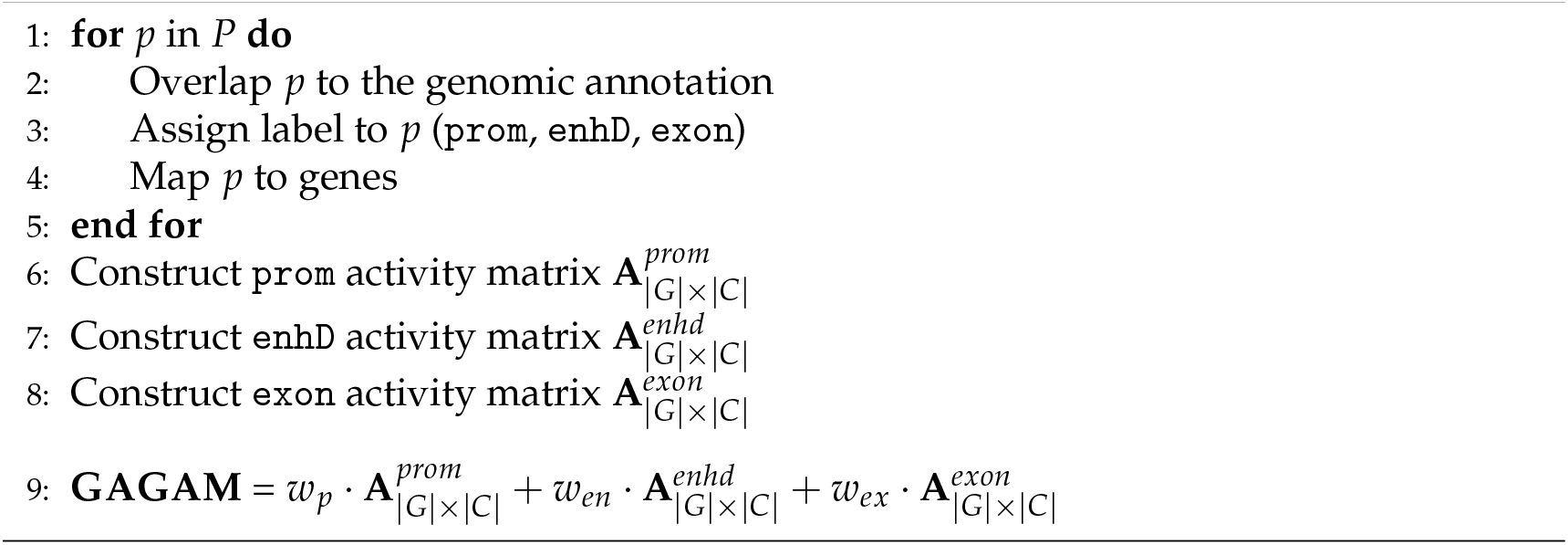

GAGAM operates solely on preprocessed scATAC-seq data organized in the form of a matrix **D**_|*P*|×|*C*|_where *P* is the set of peaks in the dataset, and *C* is the set of available cells. As illustrated in Algorithm 1 (lines 1 to 5), GAGAM utilizes the UCSC Genome Browser [20] to obtain genomic annotations, labeling all peaks *p* ∈ *P* overlapping with regions of interest (i.e., promoters, genes’ exons, and enhancers). This assigns labels prom, exon, enhD to peaks, linking accessible peaks to their biological functions. The original dataset **D**_|*P*|×|*C*|_is then split into three subsets based on the three sets of labeled peaks: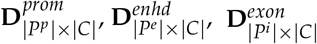. These three matrices are further processed to obtain label-specific gene activity matrices (Algorithm 1, lines 6 to 8), denoted as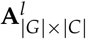 where *G* is the set of genes with at least one mapped peak and *l* ∈ *prom, enhd, exon*. Interested readers may refer to [10] [11] for a detailed description of the GAGAM model.

The current GAGAM model, while implementing cis-regulatory elements to calculate activity scores, solely considers their accessibility. Although a good indicator for assessing involvement in transcription, it overlooks the interaction with TFs. scATAC-seq data, however, offers ways to investigate TF interactions. Identifying DNA sequences where TFs can bind, known as Transcription Factor Binding Motifs (TFBMs), is feasible. However,these motifs, i.e., short sequences of a maximum of a dozen base pairs (bp), present limitations compared to the hundreds of bp in peaks [5]. Additionally, a motif does not guarantee TF binding, as the specific TF must be expressed and transcribed by cells to be actively involved in regulation. When a TF is bound to DNA, the region becomes inaccessible to the Tn5 Transposase used in scATAC-seq experiments, leaving a detectable footprint in the signal [14]. However, due to the sparsity of single-cell data, TF footprints are not measurable for each region with a specific TFBM but require studying the average signal from all motif instances.

Various types of information are available to study TFs in single-cell experiments, but looking at only one aspect has limitations. This work proposes a preliminary assessment of the correlation between TFBM enrichment, TF footprint signal, and actual TF expression. Understanding the intricate dynamics of TF contribution is crucial for proper modeling in GAGAM. The analysis examines motifs in promoter and enhancer regions independently, as this separation is central to GAGAM, and different TF interactions occur in these functional regions.

## 3. Materials and Methods

### 3.1 Dataset

This work requires a multi-omic dataset to allow a direct comparison between the epigenetic information (from scATAC-seq) and the gene expression (from scRNA-seq). The dataset of choice is an open-access dataset from the 10X Genomics platform, consisting of 10,691 cells from adult murine peripheral blood mononuclear cell (PBMC) [21]. The scATAC-seq part of the dataset has a total of 115,179 peaks as features, while the scRNA-seq part has 36,601 genes. The tools employed to process and elaborate the data are GAGAM (the focus of this paper, accessible from [10]) and Seurat [1]. The latter is one of the most well-known and highly-utilized single-cell pipelines. It allows the processing of the datasets, which is beneficial since it supports a data structure tailored to contain the results of different epigenetic analyses. Moreover, Seurat provides a dataset integration approach to label the cells with known cell-type labels by employing an external reference dataset. In this way, the cells are divided into cell-type clusters representing the ground truth of the following analyses.

### 3.2 Aggregated Cells

Before starting with the actual meta-analysis, it is worth noting that scRNA-seq and scATAC-seq detect only a tiny fraction of the actual signal from each cell (around 10-45% for scRNA-seq and only 1-10% for scATAC-seq [19]). This translates into considerable sparsity for the data. For each cell, the dataset contains several zero entries that could be false negatives [22]. This characteristic introduces noise when trying to correlate accessibility and expression. For this reason, this work explores the idea of performing the analysis based on the concept of *aggregated cell* behavior. Specifically, it aggregates cells from the same cell types obtained from the Seurat integration, representing the average over groups of cells instead of single cells. In this way, this work computes the correlation not on the single cells *c* ∈ *C* (where *C* is the set of cells of the dataset) but on the aggregated cells *ct* ∈ *CT* (where *CT* is the set of cell-types) representing the average behavior over the cell types.

### 3.3 TF motifs and motif enrichment analysis

TFBMs are DNA sequences that bind to transcription factors. Known TFBMs are curated in JASPAR [23], an open-access database storing manually curated transcription factors binding profiles across multiple species of eukaryotes. These TFBMs are not exact sequences of nucleotides since there is a natural redundancy of sequences recognized by the TFs. Therefore, the motif information is stored in a Position Frequencies Matrix (PFM), representing the probability of finding a specific base for each nucleotide in the binding region. Given that the motifs are short sequences (6-12 bp), it is likely to find redundant and non-relevant matches when searching for DNA sequences matching the motif. Therefore, instead of searching for all possible matches inside accessible regions, it is more common to implement a motif enrichment analysis. In other words, this type of analysis identifies how much a known motif is over- or under-represented in a cell’s accessible regions [14]. To do so, this work employs ChromVAR [24], an R tool for analyzing sparse chromatin accessibility data, providing reliable motif enrichment functions. The function requires the motif from JASPAR and calculates the enrichment scores for each of the *m* ∈ *M* motif inside each cell, obtaining a final matrix of **ME** _|*M*|×|*C*|_ . The set of *M* motifs in this work does not comprehend the total 632 human motifs provided by JASPAR. It narrows it to the motifs whose corresponding TFs are also expressed in the dataset since the focus is the correlation of the motif information with the TFs expression. However, instead of performing, as commonly done, this analysis on the dataset as a whole, in this work, we are interested in investigating the differences between promoter and enhancer regions. Therefore, given the two scATAC-seq sub-matrices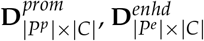, the motif enrichment calculation is performed on them separately, meaning it will capture the over- or under-representation of the motif specifically in promoter and enhancer regions. This calculation results in two matrices **ME**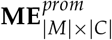and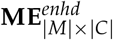, subsequently transformed into their aggregated form 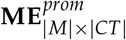 and 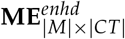 that this work analyzes separately.

### 3.4 TF footprints

TFBMs are good indicators for inferring the interaction between DNA and TFs. However, they give information only on the possible binding locations but do not capture the actual binding events. Indeed, of the millions of motifs detected on the DNA, only a small portion are actual binding regions, and even less will be relevant in a certain cell type. Fortunately, scATAC-seq data can help investigate these binding events through the TF footprint analysis. In scATAC-seq sequencing experiments, when a TF is bound to the DNA, it protects that region from sequencing, while the DNA bases immediately adjacent to TF binding are accessible, leaving, in this way, a sort of footprint in the signal. This footprint appears as a low signal from the center of the TFBM and a stronger signal from its immediate flanking regions. Ideally, the footprint would be detectable for each cell and location, but it would require a much higher sequencing depth than the technology provides. Therefore, the signal from all the motif occurrences is aggregated. Specifically, this work calculates for each motif in each cell the aggregated sequencing signal at the motif and its surrounding ±250 bp, as shown in Algorithm 2. The result is a list of | *M*| vectors**FP**_*m*_ = [ *fp*_1_, …, *f p*_*n*_] with *n* = 500+ length of the *i*^*th*^ motif, where each element represents the average normalized bias-corrected insertion signal for all the base pairs surrounding all the occurrences of the accessible motif inside a cell. This calculation is lengthy and computationally heavy for all the motifs. From it, it is possible to define the footprint score of the motif in the cells as the average signal from the motif flanking regions defined as ±50 bp from the motif.

Again, this work differentiates the signal from promoter and enhancer regions. This calculation is performed on the two matrices separately, obtaining 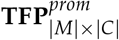 and 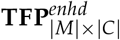 where the matrices elements are the footprint scores of motif *m*∈ *M* in cell *c* ∈ *C*. Also in this case, this work aggregates cells based on their cell-type labels, resulting in the final Matrices 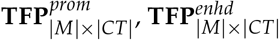where *CT* is the list of cell-types.

### 3.5 Correlation with expression

As previously highlighted, this study aims not only to explore TFs through their motif accessibility but, more significantly, to comprehend their correlation with expression levels. Before delving into this analysis, a brief overview of the gene expression matrix formalism is necessary. The considered multi-omic dataset provides a gene expression matrix **E** *G C*, encompassing 29,372 genes detected in the experiment. The matrix is then narrowed to the set *M* of genes associated with TFss, resulting in a sub-matrix **E**|*M*|×|*C*|. Subsequently, as discussed in Section 3.2, the expression values are averaged across cell types, yielding the final **E** |*G*|×|*CT*| matrix. This matrix serves as the foundation for subsequent comparisons. The correlation study follows the approach employed in [13], where the Pearson correlation between expression and motif information is calculated for each gene in the aggregated cells. These correlations are then presented through scatter plots to illustrate the general correlation within each cell type visually. The investigation covers motif enrichment versus expression and TF footprint scores versus expression, consistently differentiating between promoter and enhancer contributions.

#### Algorithm 2

Footprint score computation

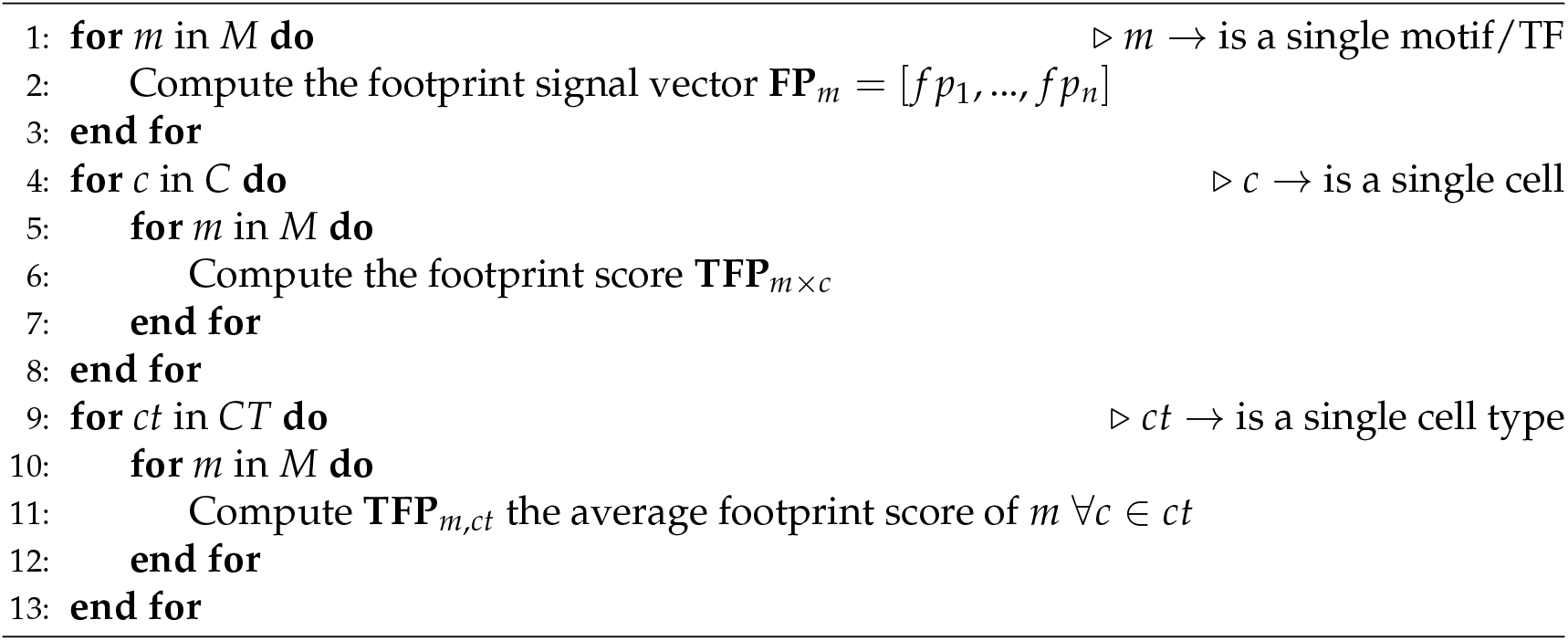

This methodology enables exploring the intricate relationship between inferable information related to TFs from accessible regions and their expression patterns.

## 4. Results

### 4.1 Enhancer regions shows more variability in motif information

When examining the motif enrichment matrices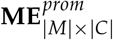And 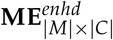 it is note-worthy that the enrichment in enhancer regions exhibits more pronounced variability compared to the enrichment in promoter regions. This contrast is clearly illustrated in Fig. 2, where the variability in TFBM enrichment is markedly higher in enhancer regions than in promoters. In studying cellular heterogeneity, this observation underscores the substantial contribution of enhancer regions in conveying relevant differences in motif accessibility. Analyzing data with higher variability is crucial for detecting distinctions between cell populations and emphasizing the significance of enhancer regions in the epigenetic context. However, the information derived from promoters should not be overlooked. Notably, from Fig. 2, it is interesting to observe that the most variable TFBMs in promoter regions belong to the FOS and JUN TF families. These families are known to aggregate and form the complex Activator Protein-1 (AP-1), which binds to promoters, regulating nuclear gene expression in T-cells. Further discussion on this TFs is provided in the subsequent section 4.2.

**Figure 1.**
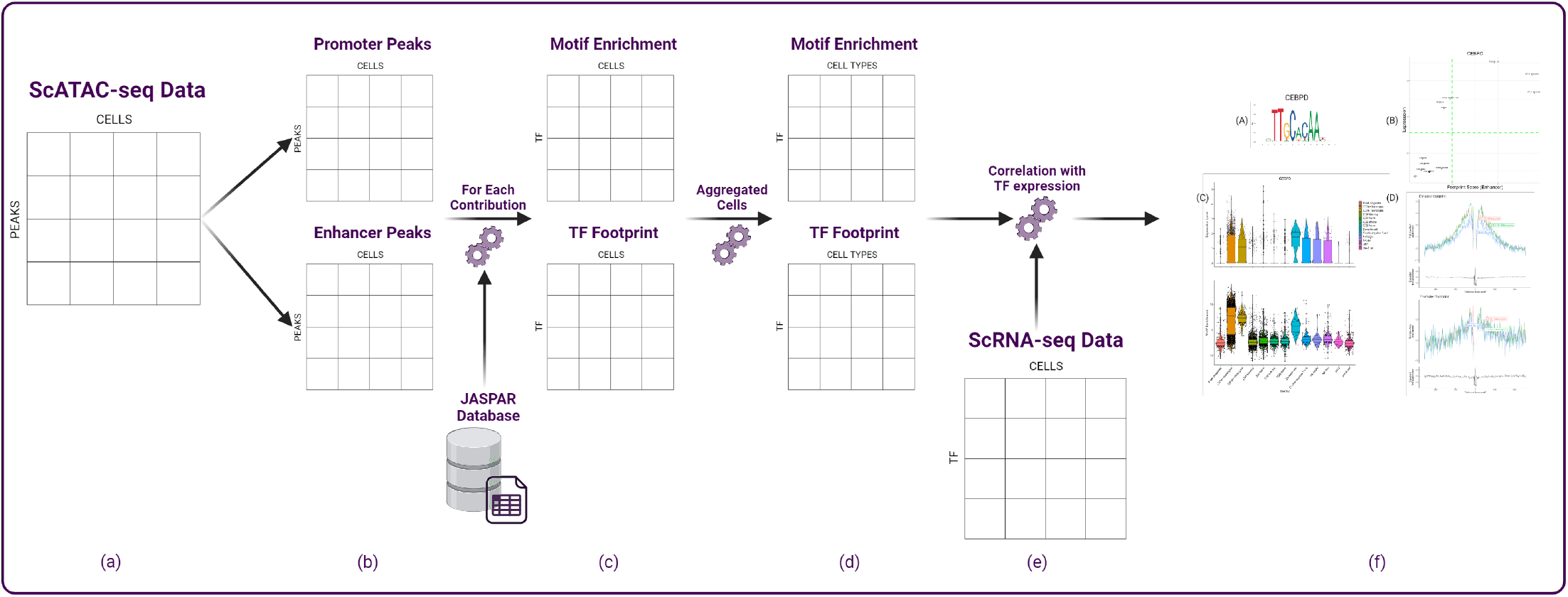
Workflow: **A** The scATAC-seq data serves as the input. **B** The data is partitioned into GAGAM contributions, resulting in promoter and enhancer matrices. **C** For each contribution, utilizing the JASPAR database, both motif enrichment and TF footprint scores are determined for all TFs expressed in the dataset. **D** Cells are aggregated based on cell-type annotations. **E** The two matrices obtained for each contribution are then compared to the expression matrix to analyze their correlation. **F** Results analysis reveals specific correlations and dynamics between TF expression and its motif information.

**Figure 2.**
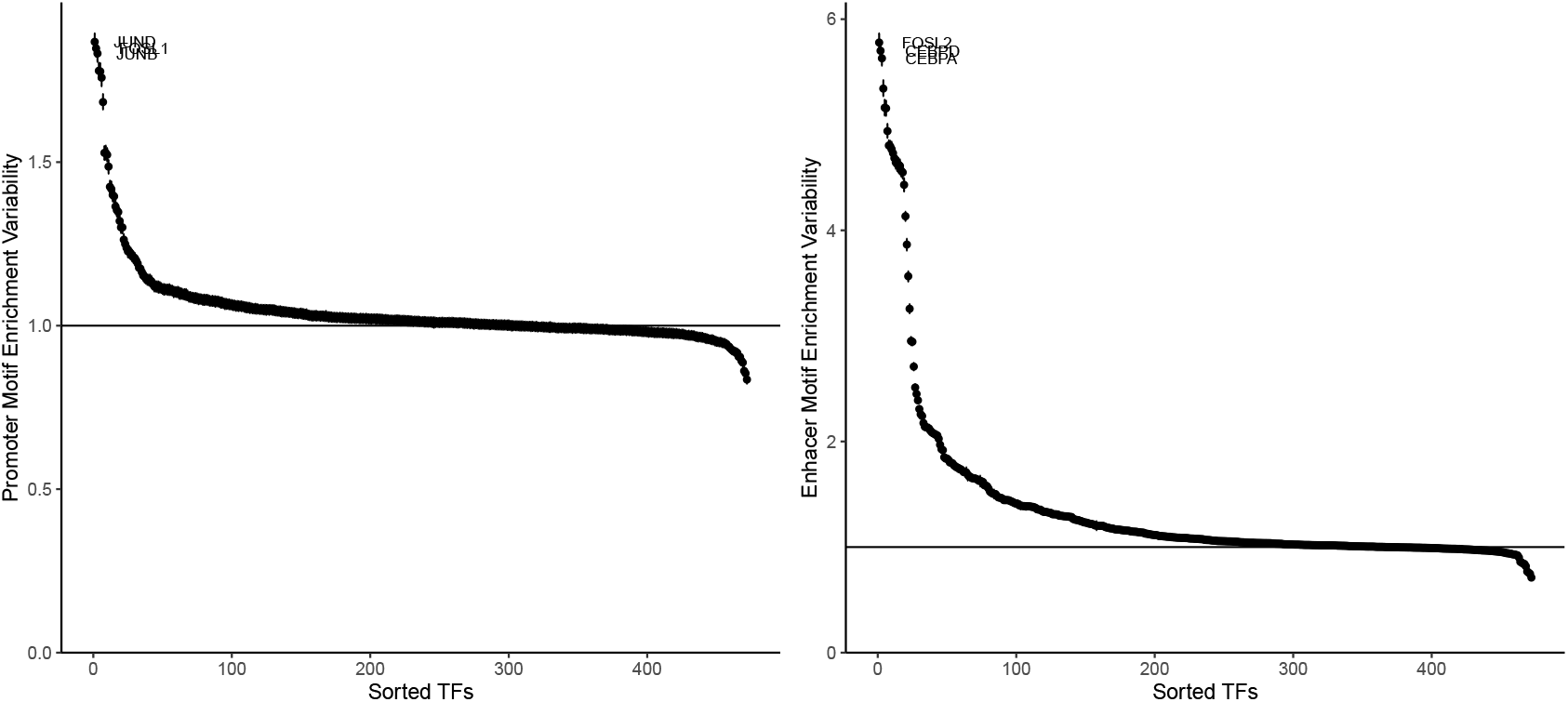
sorted TFs variability for promoter motif enrichment (left) and enhancer motif enrichment (right). For the promoter motif enrichment, the top TFs belong to FOS and JUN TF families known to bind to promoter regions.

These distinctions are illustrated in Fig. 3, where each black dot in the violin plots represents the motif enrichment of a TF for each cell type. Notably, enhancer regions exhibit only a few TFBMs with noteworthy enrichment, underscoring their significance in each cell type. It is well-established that only a subset of TFs influences gene regulation in a given cell type, especially when regulating cell-type-specific gene pathways [25]. Thus, identifying a limited number of highly enriched TFBMs per cell type aligns with expectations. Consequently, enhancer regions demonstrate a higher sensitivity to cell type-specific motif accessibility than promoter regions. This observation is consistent with findings in [13], emphasizing the substantial contribution of enhancers to the epigenetic signal in scATAC-seq and their more significant variability in correlation with expression. Once again, these results underscore the importance of modeling promoter and enhancer contributions differently, highlighting the limitations of approaches that solely focus on promoters and overlook valuable information.

**Figure 3.**
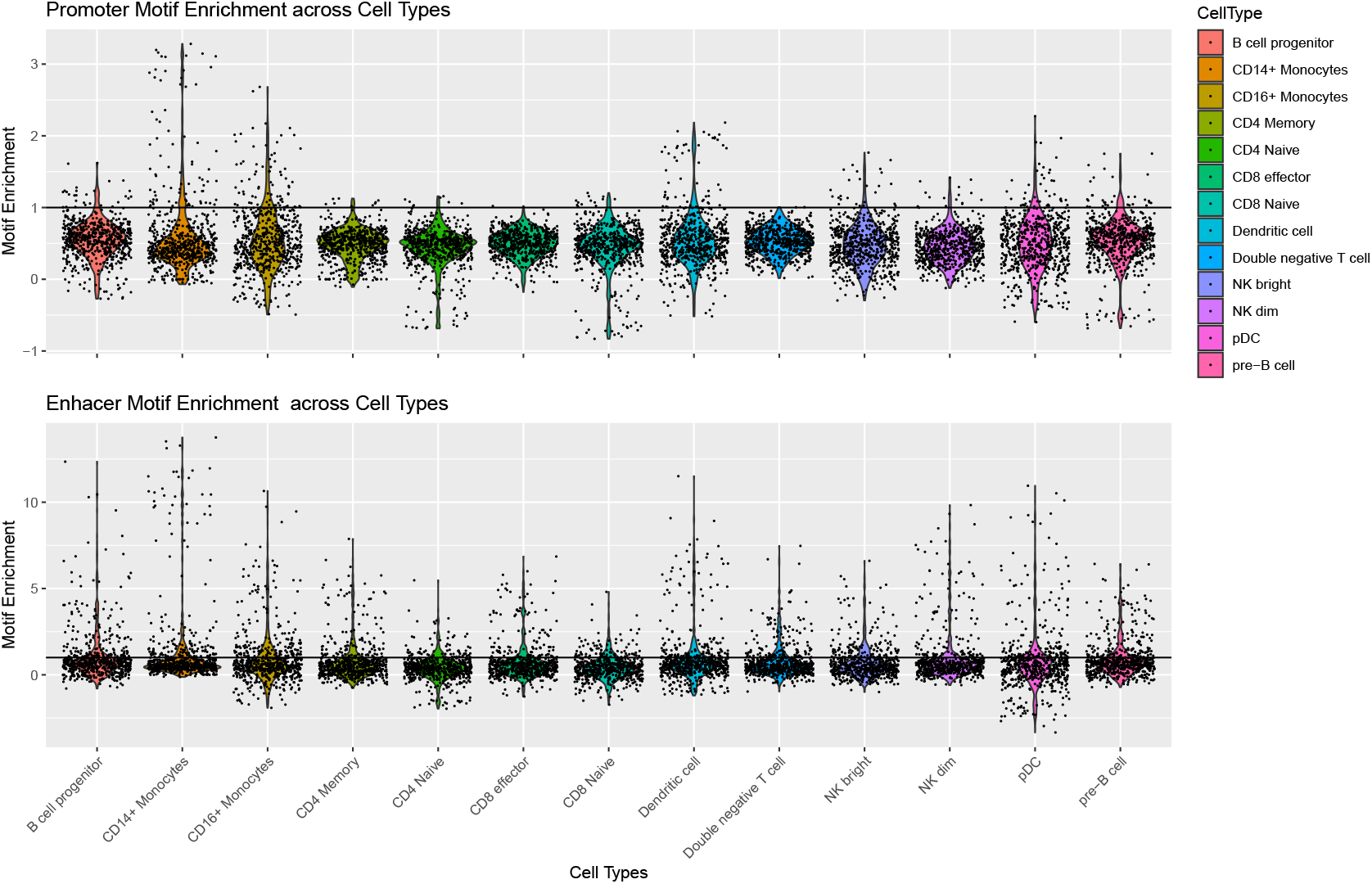
TFs motif enrichment for each cell type. Each black dot represents a TF.

### 4.2 General correlation with expression is low

Having explored the distinctions between enhancer and promoter regions, this section delves into their correlation with the actual expression of TFs. In contrast to findings in [13], interpreting the correlation between TFs expression and their accessibility information is less straightforward. The general Pearson correlation scores are relatively low. Specifically, for the expression-TFBM enrichment correlation in both promoter and enhancer regions, only around 15% of TFs exhibit a correlation higher than 0.5 (with significant p-values). Conversely, examining the expression-TF footprint correlation, 19% of TFs show a correlation higher than 0.5 (with significant p-values) in enhancer regions, which increases to 42% in promoter regions. This overall low correlation is expected, considering that, as discussed earlier, only a subset of TFs will be relevant in a particular cell type, resulting in coherent and correlated motif information for only some of them.

This observation becomes more evident when referring to Fig.4. Each point on these plots represents a single TF for each cell type. Notably, many TFs are clustered in the bottom left corner of the motif enrichment plots, indicating low expression and motif enrichment. The interesting aspect lies in the TFs characterizing each cell type, identifiable in the top right section of the plots. Here, TFs that are relevant from both expression and motif perspectives are found. Notably, these TFs tend to be cell type-specific, emphasizing that, for each cell type, a distinct subset of TFs is captured in this quadrant, aligning with the previous comments on the relevance of specific TFs for different cell types.

**Figure 4.**
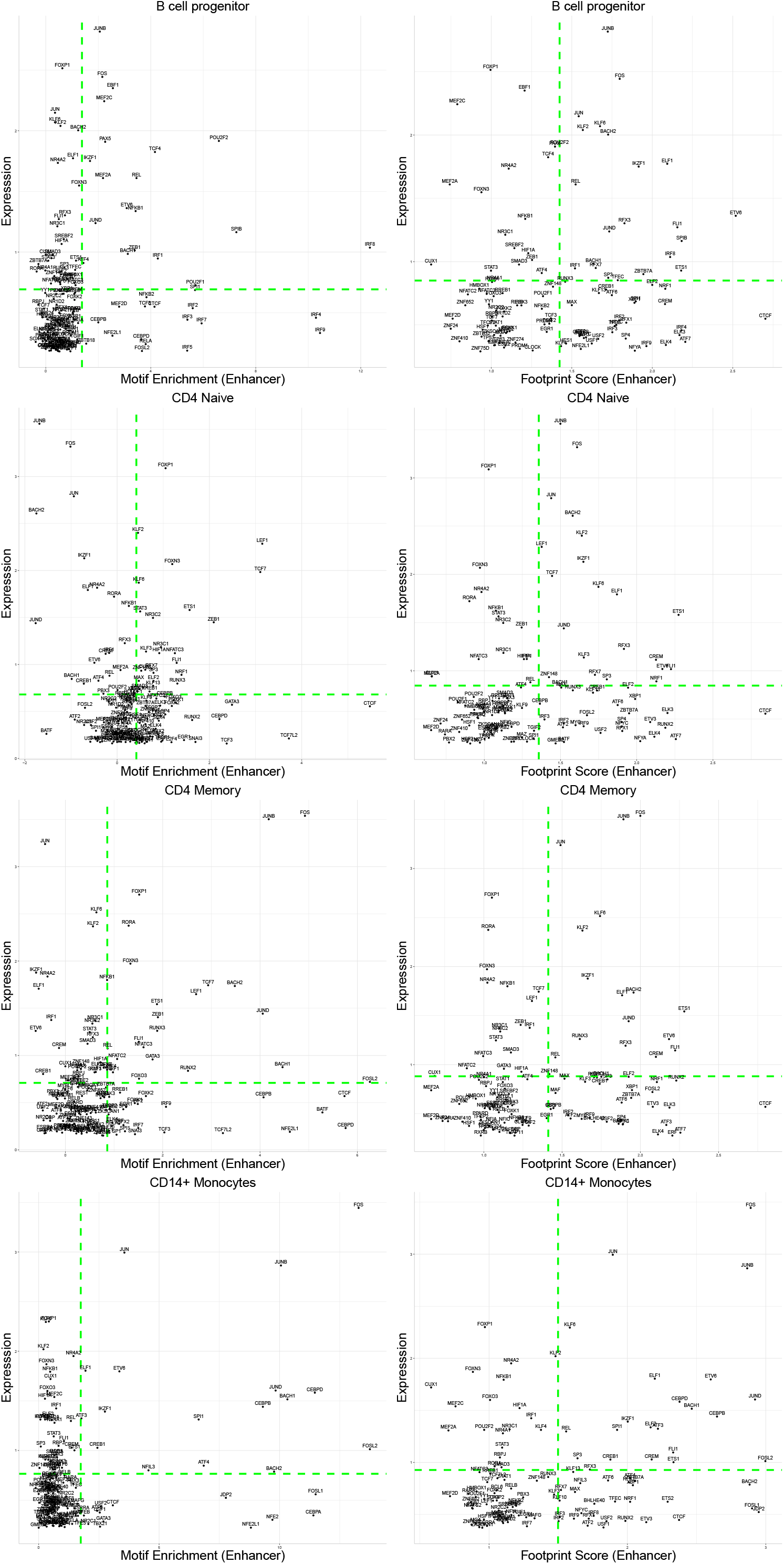
Scatter plots for the Expression-Motif Enrichment and Expression-TF footprint correlation.

Furthermore, these cell-type-specific TFs show consistency between the two motif levels. Examples include BACH1, CEBPB, and CEBPD, which are characteristic for CD14+ Monocytes in both motif enrichment and footprint score. This consistent dual information is crucial for identifying and studying cell-type-specific TF mechanisms and regulation, not apparent when solely considering expression levels. CTCF, for instance, appears to have a strong signal in all cell types despite low expression. This characteristic is discernible only when examining enhancers, highlighting a specific correlation between CTCF and enhancer regions. This aligns with its known role in DNA bending, facilitating interaction between promoters and enhancer regions [26].

Lastly, beyond these general observations, investigating differences between differentiated cell types, such as Naive and Memory CD4+ T-cells, could unveil specific TF motif patterns involved in differentiation or proliferation processes. This aspect is explored in the following section, examining various types of correlation for selected TFs.

### 4.3 Motif information highlights differences in AP-1 subunits

The AP-1 TF is a dimeric transcription factor composed of two subunits, typically belonging to the Fos and Jun TF families. It is a key regulator in processes such as cell proliferation, differentiation, and apoptosis [27] [28]. Numerous studies highlight its impact on T-cell activation, the initial step driving the differentiation and proliferation of Naive T-cells into various specialized T-cell subsets [29] [30]. As evident from the scatter plots (Fig. 4), both FOS and JUNB consistently occupy the top region for all cell types. This trend is further elucidated in Fig. 5 and Supplementary Fig. 1, which present various details for FOS and JUNB, respectively. The violin plots depict the expression (B, top) and motif enrichment (B, bottom) of the TF in each cell type. While the gene expression appears relatively consistent across cell types, there is a distinct separation in motif enrichment.

**Figure 5.**
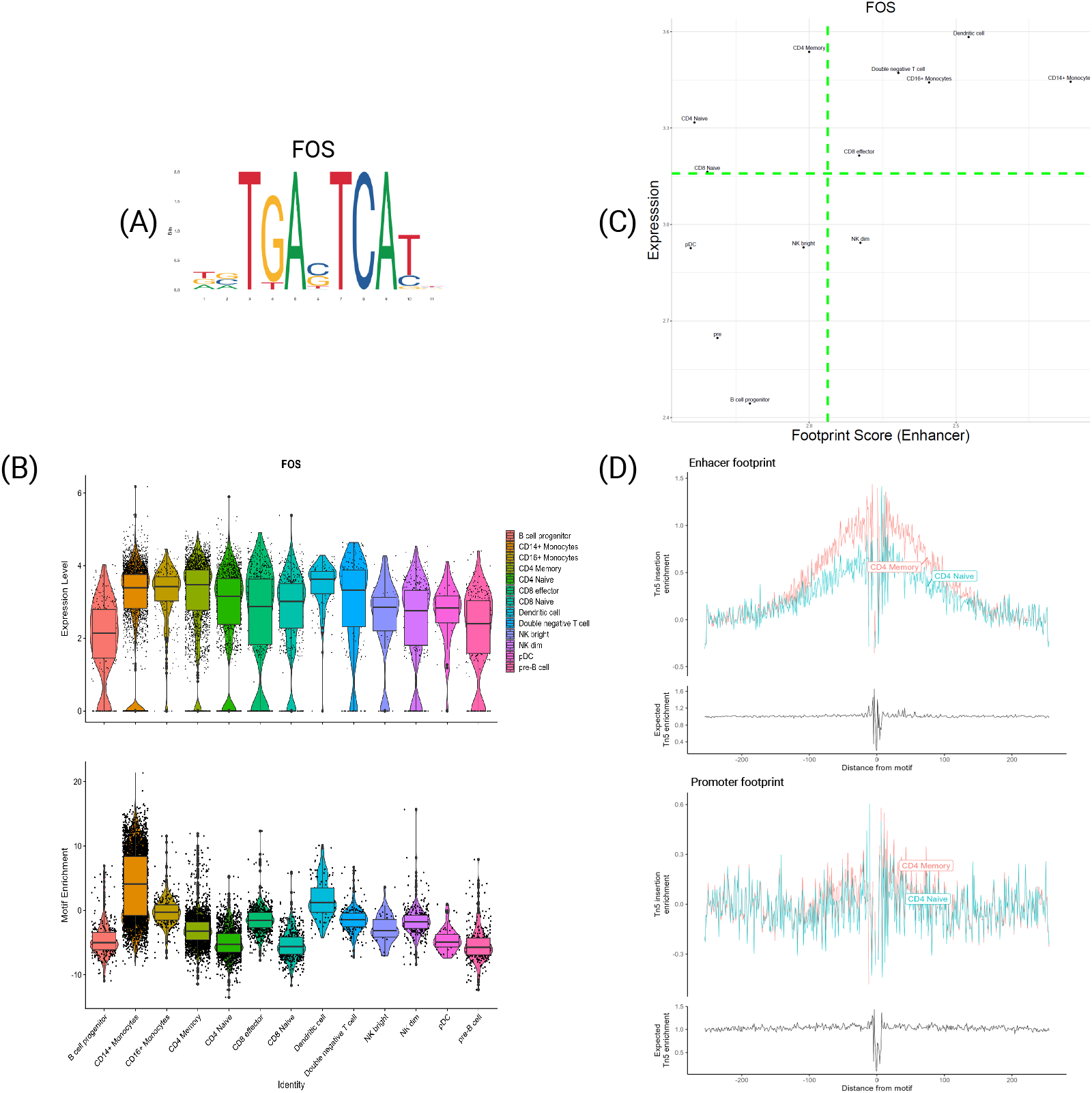
Transcription Factor FOS. **A** PFM visualization for FOS’s motif MA0476.1. **B** Violin plot of expression (top) and motif enrichment (bottom), for all cell types. **C** Scatter plot of footprint score-expression of FOS for all cell types. **D** Tn5 insertion plots for Memory and Naive CD4 T-cells.

Crucially, CD4 Memory and CD8 Effector T-cells exhibit significantly higher enrichment than their respective Naive T-cells, aligning with the discussed activation mechanism. This pattern is reinforced by the TF footprints, as illustrated in Fig. 5-C, representing the expression-TF footprint of FOS for each cell type. Once again, Naive T-cells exhibit noticeably lower footprint scores than their counterparts. This distinction becomes more apparent in the plots in Fig. 5-D, showing the average signal around the motifs. For clarity,only CD4 Naive and Memory cells are depicted, demonstrating the stronger footprint signal for Memory T-cells. Importantly, this variation is evident only for enhancers and not promoters, underscoring the notable differences between signals from functional genomic regions.

A similar pattern is observed for the gene BATF. As depicted in Supplementary Fig. 2, the expression is uniformly low across all cell types. However, both motif enrichment and footprint scores exhibit variations between naive and memory cells, especially noticeable between CD8 Naive and CD8 Effector T-cells. BATF is recognized for its involvement in the functional development of CD8 T-cells [31,32], once again emphasizing how the dynamic of TFs expression alone may not suffice to discern specific changes between cell types.

In conclusion, the FOS and JUNB TF, subunits of AP-1, exhibit a significant correlation with certain cell types in their motif information despite minimal variability at the transcriptional level. This result, coupled with observations on BATF, underscores that TFs expression alone may not be adequate to identify crucial differences in specific cellular processes; it must be complemented by an examination at the epigenetic level for a more comprehensive understanding.

### 4.4 Specific TFs shows differences only at the expression levels

FOS and JUNB TFs are not the only ones with differences in expression and motif information behavior. Indeed, the TF BACH2 has a likewise interesting behavior. BACH2 is a known regulator in the B-cells development [33], by orchestrating the early specification and commitment of B-cell progenitors [34]. However, Fig. 6 shows a peculiar behavior in its motif information. Looking at Fig. 6-B, it is evident that there is a gene expression differentiation between cell types, particularly between B-cell progenitors and pre-B-cells, being overexpressed in the latter. However, the motif enrichment in these two cell types is pretty much identical, implying that only the TF expression drives this differentiation, an opposite behavior to FOS and JUNB. Furthermore, the footprint plots in Fig. 6-D show no difference between the two cell types.

**Figure 6.**
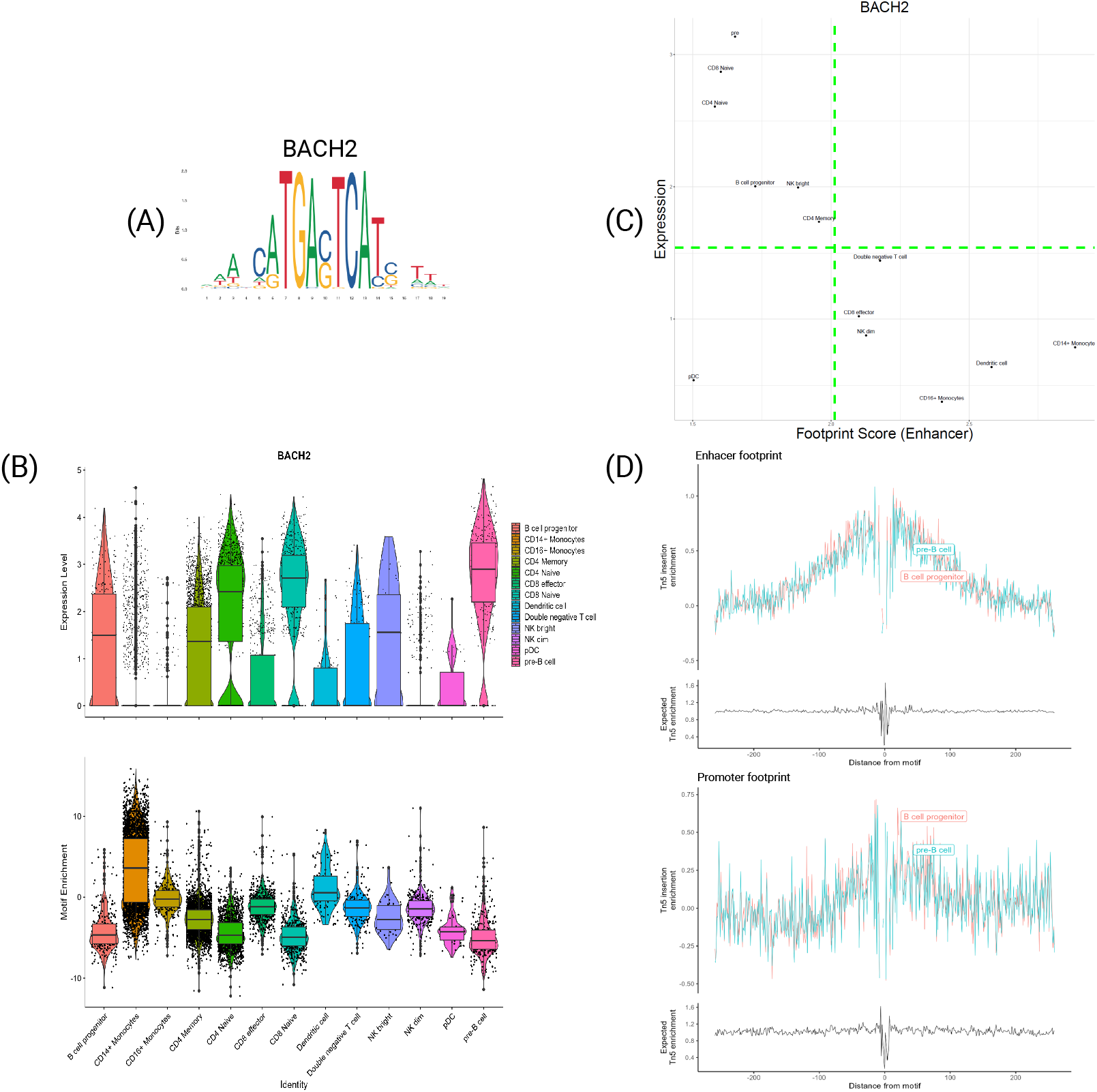
Transcription Factor BACH2. **A** PFM visualization for BACH2’s motif MA1101.2. **B** Violin plot of expression (top) and motif enrichment (bottom), for all cell types. **C** Scatter plot of footprint score-expression of BACH2 for all cell types. **D** Tn5 insertion plots for B-cell progenitors and pre-B-cells.

Furthermore, the motif enrichment in the other cell types seems to have an inverse trend to the expression. Indeed, cell types with the highest expression (such as Naive T-cells and pre-B-cells) display the lowest enrichment. This inverse correlation is even more evident from Fig.6-C, showing how cell types with lower expression have higher footprint scores and vice versa. This peculiar behavior seems counterintuitive since one would expect not to see a higher expression if the motif is that much accessible. However, from the literature, BACH2 is highly characterized as a repressor TF [35], which regulates

B-cells by suppressing specific genes related to the myeloid program. Hence, the observed inverse correlation for this gene may indicate a distinctive repressive dynamic. In cells where BATCH2 is active, the motif of this gene becomes generally less accessible but more specific, fine-tuning its repression mechanism and subsequently leading to an increase in its expression. BACH2 exhibits a unique connection between its expression and motif information, serving as an intriguing indicator to discern repressive dynamics from the more common enhancing dynamics. This observation is of utmost importance, as an effective transcriptional regulation model should accurately represent these two markedly different dynamics. Currently, there remains a need for consensus or comprehensive information to distinguish the intricacies of silencing processes.

### 4.5 TFs characterize cell types at both expression and motif information level

In this concluding section, it is pertinent to showcase genes that exhibit coherent and cell-type-specific correlations between their expression and motif information. For instance, the transcription factor CEBPD, depicted in Fig. 7, is recognized for its role in the inflammatory response of monocytes [36–38]. This functional role is reflected in its expression and motif information, as illustrated in Fig. 7-B. CEBPD is selectively expressed in monocytes and dendritic cells while significantly over-enriched in these cell types. This observation is reinforced by the footprint score, where the scatter plot indicates that these subtypes are situated in the top-right corner, signifying that CEBPD plays a regulatory role in these cells at both the transcriptomic and epigenomic levels.

**Figure 7.**
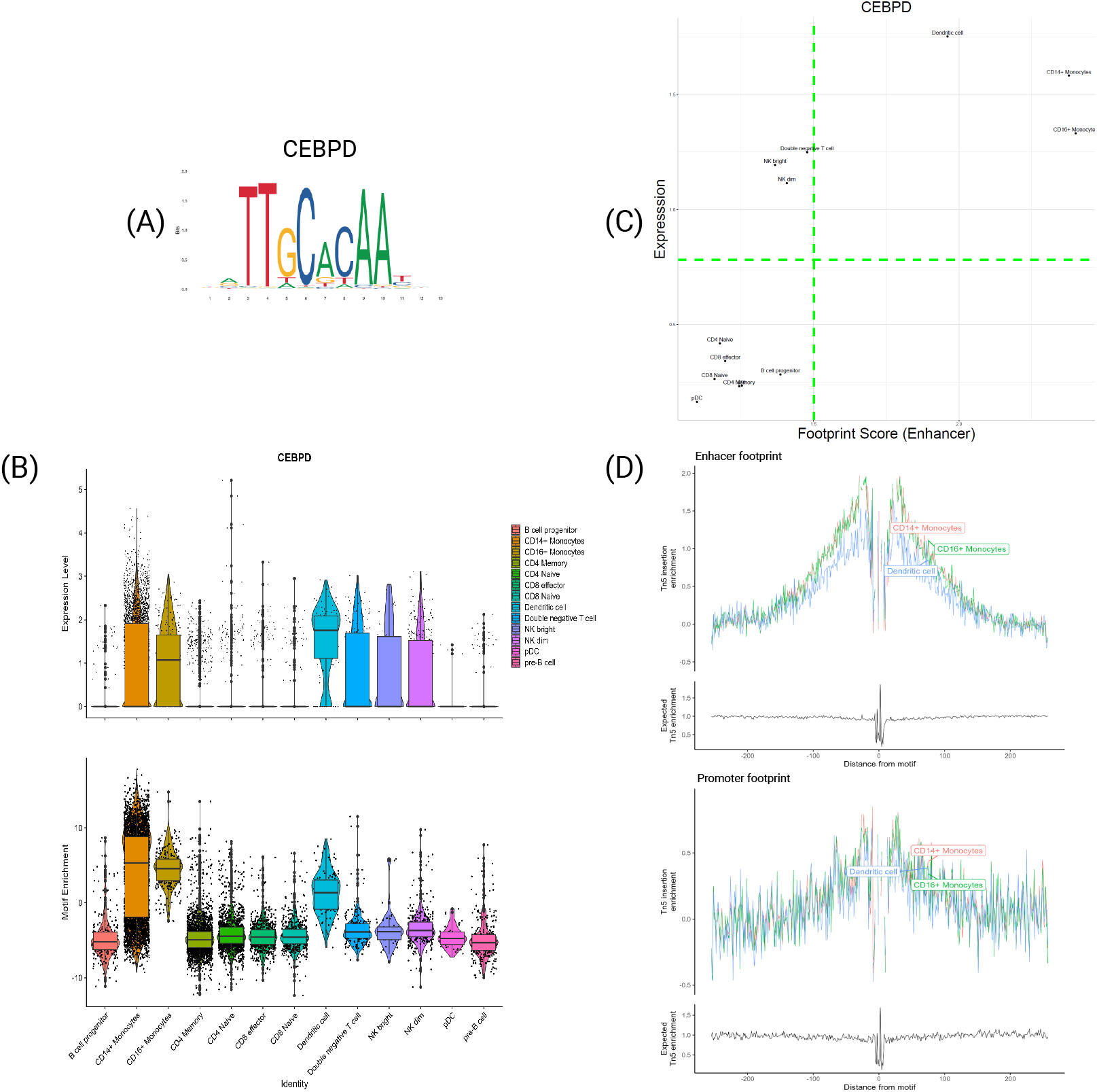
Transcription Factor CEBPD. **A** PFM visualization for CEBPD’s motif MA0836.2. **B** Violin plot of expression (top) and motif enrichment (bottom), for all cell types. **C** Scatter plot of footprint score-expression of CEBPD for all cell types. **D** Tn5 insertion plots for CD14+ and CD16+ Monocytes and Dendritic cells.

Similarly, Fig. 8 presents the findings for the TF POU2F2. This TF holds considerable significance in B-cells, particularly in motif enrichment, as it exhibits the highest motif enrichment signal among those investigated. This observation aligns with the footprint in Fig. 8-D, showcasing a prominent signal at the flanking regions for both subtypes. Once again, the scatter plot illustrates that B-cells consistently reside in the top-right corner, indicating high expression and footprint scores, while other subtypes occupy the opposite bottom-left corner.

**Figure 8.**
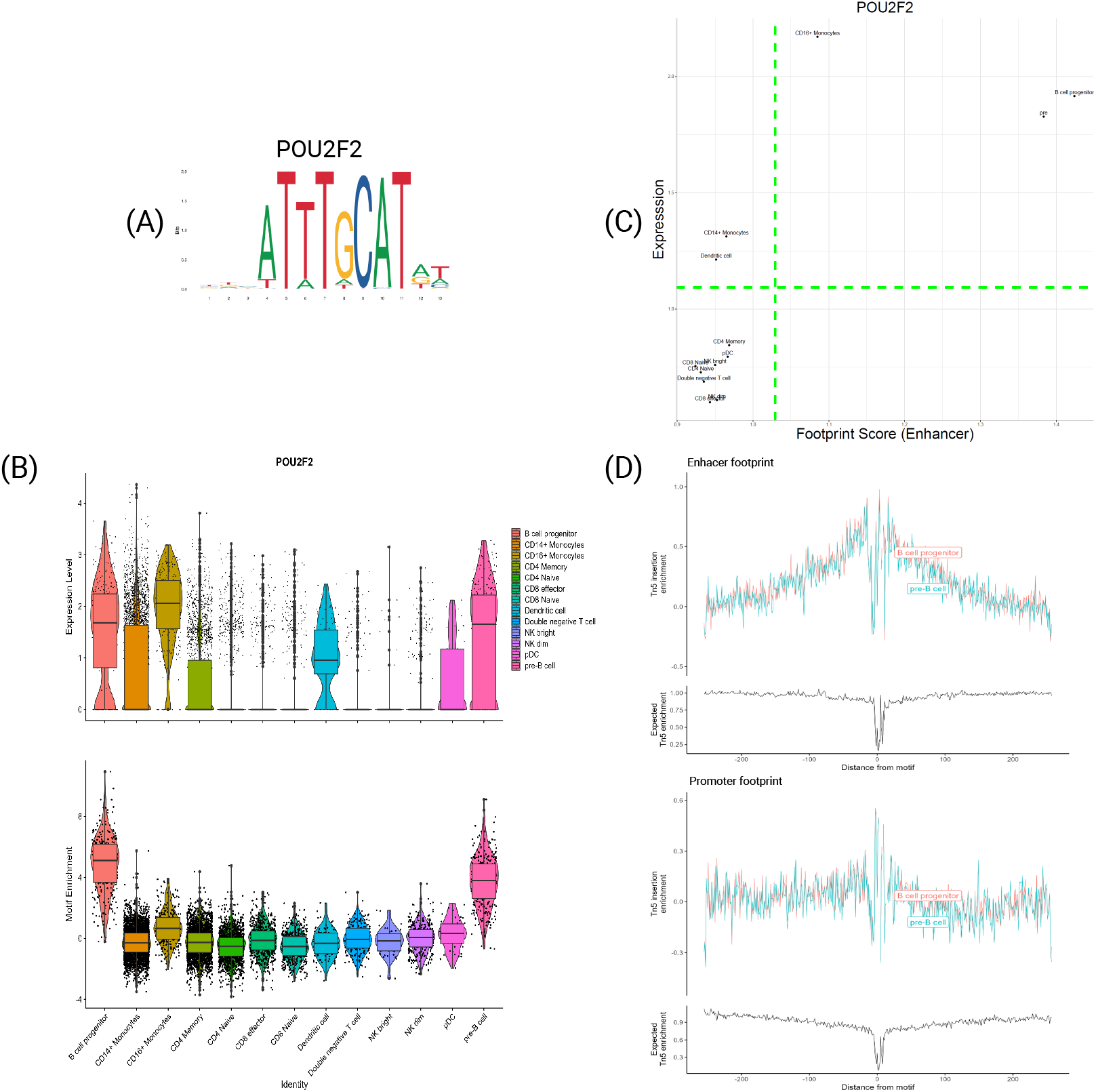
Transcription Factor POU2F2. **A** PFM visualization for POU2F2’s motif MA0507.1. **B** Violin plot of expression (top) and motif enrichment (bottom), for all cell types. **C** Scatter plot of footprint score-expression of POU2F2 for all cell types. **D** Tn5 insertion plots for B-cell progenitors and pre-B-cells.

Similar observations apply to the TFs TCF7 and LEF1 (see Supplementary Fig. 3 and 4). Both are characteristic of T-cells and demonstrate coherence between expression and motif information. Notably, there is a discernible difference in CD8 effector cells, which exhibit a weaker correlation than other T-cell subtypes. This discrepancy may highlight a specific dynamic of these TFs in those subtypes, which could be crucial in distinct biological processes.

In conclusion, what ties these genes together is their specific relevance in distinct subtypes at both the expression and motif information levels. Therefore, modeling their impact on transcriptional regulation is essential, as they likely serve as specialized regulators characterizing various biological processes.

## 5. Conclusions

This work presents a comprehensive analysis of the correlation between the motif information obtainable from scATAC-seq data and the expression of the TFs themself. This analysis is crucial for understanding transcriptional regulation, which the TFs are a crucial part of. The increasing power of multi-omic sequencing technologies assists in this, simultaneously allowing the investigation of the expression and DNA accessibility of relevant regions related to transcriptional regulation. Specifically, this work investigates the motif presence in accessible enhancer and promoter regions distinctively. Two types of information are considered: the motif enrichment, representing how much a motif is over-represented in determined regions, and the TF footprint scores, representing the signal of a TF binding event. This analysis brought some interesting results. First, there is a remarkable difference between the signal from enhancers and promoters, with the first showing a more significant variability between cell types and highlighting different types of TFs. These differences show the importance of a distinct analysis of the two types of functional regions, which need to be studied separately to properly understand the intricacies of transcriptional regulation.

However, the correlation between the motif information and the expression is low, whatever contribution one considers. This result is partially expected since only a small subset of the TFs is cell-type specific and, consequentially, is coherent between the omic levels. However, the reported results highlight differential behaviors of specific TFs between certain cell types. The exciting part is that the results highlighted different correlation patterns in the motif information, indicating the necessity of modeling their impact on transcription in specific manners.

This work focused on the TFs by themselves, looking at the different information inferred by the multi-omic data. However, this is only the first step in modeling the transcriptional regulation. Future work will aim to understand the correlation between the TFs and their putative target genes, trying to understand how the motif information from scATAC-seq data can influence gene expression. Moreover, it will be relevant to practically model this correlation in an extension of the gene activity concept, specifically GAGAM, which will not only investigate the general DNA accessibility but will consider a higher level of information to model the transcriptional regulation correctly.

## Author Contributions

Conceptualization, L.M.; methodology, L.M.; software, L.M.; validation, L.M.; formal analysis, L.M. and R.B.; investigation, L.M. and R.B.; resources, L.M.; data curation, L.M.;writing—original draft preparation, L.M., R.B. and S.D.C.; writing—review and editing, L.M., R.B.,A.S. and S.D.C.; visualization, L.M., R.B., A.S. and S.D.C.; supervision, A.S. and S.D.; All authors have read and agreed to the published version of the manuscript.

## Institutional Review Board Statement

Not applicable

## Informed Consent Statement

Not applicable

## Data Availability Statement

- The data presented in this study are openly available in NCBI GENE Expression Omnibus (GEO) with accession number GSE96769, and from the freely available 10XGenomic platform at https://github.com/smilies-polito/GAGAM Accessed on 29 December 2022.
- All the code employed for this work is available at https://github.com/smilies-polito/MAGA, including all the supplementary material and figures, accessible in zenodo with the DOI https://doi.org/10.5281/zenodo.10517230

